# A Generalized *Adder* mechanism for Cell Size Homeostasis: Implications for Stochastic Dynamics of Clonal Proliferation

**DOI:** 10.1101/2024.09.13.612972

**Authors:** César Nieto, César Augusto Vargas-García, Abhyudai Singh

**Affiliations:** Department of Electrical and Computer Engineering, University of Delaware. Newark, DE 19716, USA; AGROSAVIA - Corporacion colombiana de investigacion agropecuaria, Mosquera, 250047, Colombia; Department of Electrical and Computer Engineering, Biomedical Engineering, Mathematical Sciences, Interdisciplinary Neuroscience Program, University of Delaware, Newark, DE 19716, USA

## Abstract

Measurements of cell size dynamics have revealed phenomenological principles by which individual cells control their size across diverse organisms. One of the emerging paradigms of cell size homeostasis is the *adder*, where the cell cycle duration is established such that the cell size increase from birth to division is independent of the newborn cell size. We provide a mechanistic formulation of the *adder* considering that cell size follows any *arbitrary non-exponential growth law*. Our results show that the main requirement to obtain an *adder* regardless of the growth law (the time derivative of cell size) is that cell cycle regulators are produced at a rate proportional to the growth law and cell division is triggered when these molecules reach a prescribed threshold level. Among the implications of this generalized adder, we investigate fluctuations in the proliferation of single-cell derived colonies. Considering exponential cell size growth, random fluctuations in clonal size show a transient increase and then eventually decay to zero over time (i.e., clonal populations become asymptotically more similar). In contrast, several forms of non-exponential cell size dynamics (with *adder-based* cell size control) yield qualitatively different results: clonal size fluctuations monotonically increase over time reaching a non-zero value. These results characterize the interplay between cell size homeostasis at the single-cell level and clonal proliferation at the population level, explaining the broad fluctuations in clonal sizes seen in barcoded human cell lines.

## Introduction

The size of cells critically impacts all aspects of their physiology regulating essential processes including gene expression regulation, nutrient metabolism and proliferation [1–7]. The factors determining cell size, and its maintenance around a desired setpoint specific to a cell type are fundamental questions that have long fascinated biologists. To achieve size homeostasis, cells must synchronize multiple processes such as chromosome replication [8], growth rate [9], cell membrane synthesis [10], organelle duplication [11], and divisome formation [12]. Remarkably, recent findings in bacterial, fungi, animal, and plant cells have uncovered common principles by which aberrations in cell size are corrected through size-dependent duration of cell -cycle stages [13–19].

One such phenomenological principle of cell size homeostasis is the adder, where for an isogenic cell population under stable growth conditions, the size added (or increment) from cell birth to division is uncorrelated with newborn size (Figure 1). Thus, regardless of how large or small a cell is born, the division timing is set to add, on average, a fixed cell size [20, 21]. This constant size increment itself depends on growth conditions and can vary considerably across cell types. For an exponential increase in cell size, an adder implies an inverse correlation between the cell cycle duration and the newborn cell size [22].

**Figure 1:**
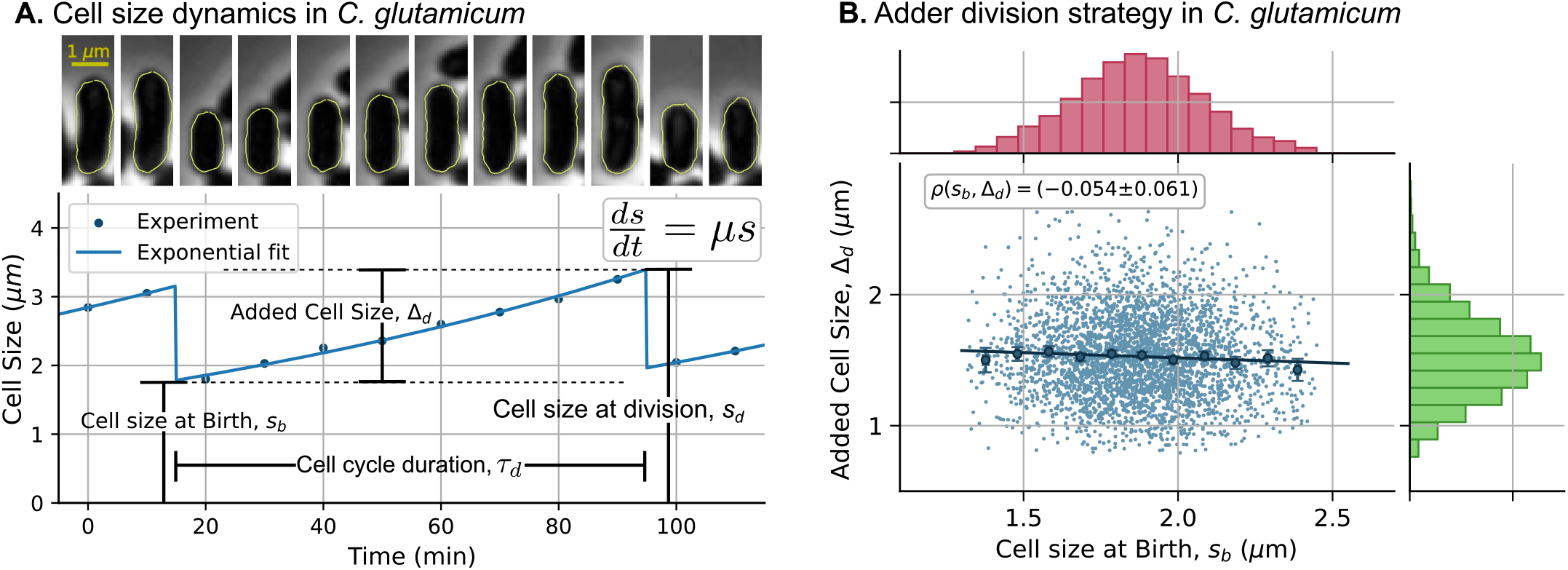
Cell size homeostasis in *C. glutamicum* occurs via the *adder* mechanism. **(A)** *Top:* Timelapse of a *C. glutamicum* cell undergoing division, with one of the newborns randomly chosen for cell size tracking. *Bottom:* The cell size (quantified by cell length) over time (dots) fits relatively well with exponential growth in cell size (solid line). Key cell cycle variables are illustrated on the plot: the growth law 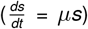, cell size at birth (*s*_*b*_), cell size at division (*s*_*d*_), added cell size from birth to division (Δ_*d*_ = *s*_*d*_ − *s*_*b*_), and cell cycle duration (*τ*_*d*_). **(B)** Data reveals adder-based cell size control, where Δ_*d*_ is uncorrelated to *s*_*b*_ across *C. glutamicum* cell cycles. The marginal plots show histograms of cell size at birth (red) and added cell size (green). Data are taken from [34], and we refer readers to this source for experimental details and image segmentation for cell size measurements. The circles on the plot correspond to the mean added size for binned newborn sizes (11 bins), with error bars representing the 95% confidence interval of the mean for each bin. The line represents the linear fit and *ρ* is the correlation coefficient between *s*_*b*_ and Δ_*d*_, with the 95% confidence interval obtained using bootstrapping.

The *adder* was discovered with the development of microfluidic imaging in evolutionary divergent bacteria such as *E. coli, B. subtilis* [23], *C. crescentus* [24], and *P. aeruginosa* [25]. Cell size analysis for organisms more complex than rod-shaped bacteria presents difficulties that have been recently solved. For example, the variable morphology of mammalian cells complicates the estimation of cell size. With advances in methods that combine measurements of cell size and cell mass, it has been observed that the *adder* principle is prevalent even in different human cell lines [26]. Some of these cell lines exhibit non-exponential growth [26, 27], which poses challenges to existing models that mainly account for the *adder* in exponentially growing cells [12, 28]. The *adder* principle has also been observed in different species, including budding yeast [29], *Dictyostelium* amoebas [30], bacterial strains that exceed exponential growth rates [31], and even flatworms that consistently increase their body area from the head after transverse splitting [32].

In this study, we investigate the minimal theoretical requirements to achieve an *adder* in cell size dynamics under any *arbitrary growth law* (the time derivative of cell size). We propose a phenomenological approach that employs a size-dependent division rate that results in an adder with an arbitrary added-size distribution. We compare this approach with an equivalent mechanistic model that involves a cell cycle regulator expressed at a rate proportional to the growth law, with cell division triggered when the regulator reaches a critical threshold level [33]. We show that, under these conditions, stochastically formulated models result in an *adder*, where the size added from cell birth to division is statistically *independent* of the initial size.

Recent studies have shown that adder-based cell size homeostasis significantly influences the stochastic dynamics of clonal populations. Specifically, clonal expansion models based on the *adder* principle produce qualitatively different predictions compared to classical time-based division models [34, 35]. Mathematical predictions and experimental observations indicate that exponentially growing cells exhibit a high degree of clonal size regulation (low stochastic fluctuations) when following the adder mechanism [34]. Motivated by these findings, we investigate how cell proliferation statistics change when cells adhere to growth laws other than exponential cell size dynamics.

Our findings indicate that exponentially growing cells that implement the *adder* exhibit clonal size fluctuations with a transient increase until the first generation, followed by a decay to zero over time. This means that cell colonies become asymptotically more homogeneous relative to each other in terms of their population size. In contrast, for non-exponential cell size dynamics, clonal size fluctuations increase monotonically over time, leading to greater heterogeneity in clone sizes as they grow. These insights are crucial for understanding noise mechanisms affecting cell lineage expansion, with significant implications for diverse fields such as cancer biology [36], embryogenesis, mutant fixation [37], and microbial ecology [38].

## Results

### A generalized *adder* mechanism

In this section, we present a generalized formalism to achieve an *adder* strategy for cell size regulation. We begin by introducing the main cell cycle variables. Figure 1 illustrates the cell size dynamics during one cell cycle of the gram-positive, rod-shaped bacterium *C. glutamicum*. Cell cycle begins with the division of the parent cell into two newborn descendants. We randomly select one of these newborns and track its size *s*(*t*) *>* 0, over time *t* until its division. For each cell cycle, we measure the cell size at birth *s*_*b*_ and at division *s*_*d*_, the cell cycle duration *τ*_*d*_, and the size added from birth to division Δ_*d*_. The relationships between these variables are fundamental for characterizing cell size homeostasis mechanisms. For instance, Figure 1B illustrates the *adder* mechanism, where the size added from birth to division is uncorrelated with the newborn size.

We assume that the cell is born at time *t* = *t*_0_ with size *s*_*b*_. Then, its cell size evolves according to the *time-varying* ordinary differential equation:

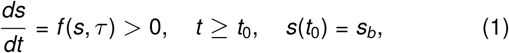

where *τ* = *t*− *t*_0_ is a timekeeper variable measuring the time spent within the cell cycle since birth. The function *f >* 0 is referred to as the *growth law* and is allowed to be an arbitrary function of the cell size and timer. Given an initial size at birth *s*(*t*_0_) = *s*_*b*_, the growth law is assumed to take a form that results in a unique solution *s*(*t*) for all *t*≥ *t*_0_. The positivity of the growth law *f >* 0 ensures a strictly monotonic increase in cell size over time until the cell undergoes division. We define another monotonically increasing function:

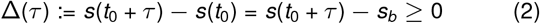

that is the size added at time *τ* after cell birth with Δ(0) = 0. Considering *τ*_*d*_ as the cell cycle duration, a random variable, one can define the added size from cell birth to division as:

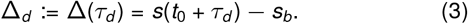

Note that since both *τ*_*d*_ and *s*_*b*_ are random variables, Δ_*d*_ is also a random variable, having different values for different cell cycles (Figure 1B). Our aim is to define a propensity for cell division that ensures two properties:

1. An *adder* for any growth law *f* where the added size Δ_*d*_ becomes independent of the newborn size *s*_*b*_. Note that while experimentally an *adder* is often characterized by Δ_*d*_ being uncorrelated to *s*_*b*_ (as in Figure 1B), here we impose a stricter condition of statistical independence.
2. The added size Δ_*d*_ follows an arbitrary given probability density function (pdf) 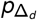, i.e, the probability that the added size Δ_*d*_ is in the interval (*y, y* + *dy*), follows the equation:

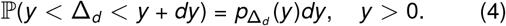

Our key result is as follows: given *s* and *τ* at any time instant in the cell cycle, a cell division event occurs with propensity *f* (*s, τ*)*h*(Δ(*τ*)), i.e., the probability of division in the next infinitesimal time interval (*t, t* + *dt* ] is *f* (*s, τ*)*h*(Δ(*τ*))*dt*, where *h* is an arbitrary positively valued function of the added size. Referring the reader to supplementary section S1 for a detailed proof, with the chosen division propensity, Δ_*d*_ becomes independent of *s*_*b*_ with pdf satisfying

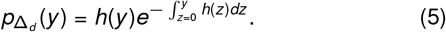

Conversely, it can be shown that choosing

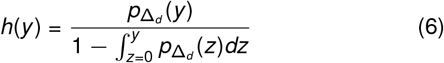

implements an *adder* for any arbitrary 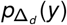 (*y*). Thus, irrespective of newborn size and growth law, the cell cycle timing is set such that the added size Δ_*d*_ is an independent and identically distributed random variable with pdf 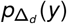 (*y*). As a simple example, the division propensity *kf* (*s, τ*) for a constant *h* = *k >* 0 results in Δ_*d*_ being an exponentially-distributed random variable with mean Δ_*d*_ = ⟨1*/k*⟩. Here, we use the notation ⟨ ⟩ to denote the operation of expected value. Similarly, for this example, the randomness of Δ_*d*_, quantified by the squared coefficient of variability

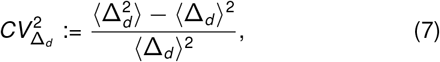

satisfies 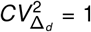

Upon cell divsion, the size and timer are reset as

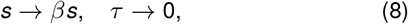

which correspond to selection of one of the daughters for further size tracking. In addition, 0 *< β <* 1 is an independent and identically distributed random variable. For symmetric division in which mother cell size is approximated halved between daughters, the mean value of *β* is ⟨*β*⟩ = 1*/*2, and its squared coefficient of variation 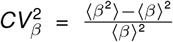 quantifies the error in size partitioning between daughters [39]. In general, it is possible to consider any arbitrary mean ⟨*β* ⟩ for non-symmetric cell size partitioning, as occurs in the case of budding yeast [40]. For ⟨*β*⟩= 1*/*2, the newborn cell size statistics (mean and coefficient of variation squared, respectively) are given by [39, 41, 42]

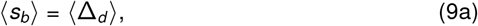

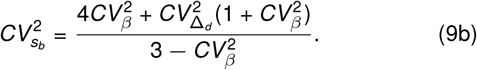

Note from (9b) that the noise in the newborn size 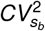 is much more sensitive to errors in cell size partitioning 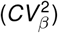 than the noise in added size 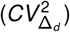. This point is more clearly seen in the limit 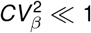 when (9b) simplifies to

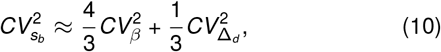

which means that *β* contributes approximately 4/3 of its noise to the newborn cell size variability while Δ_*d*_ contributes only 1/3 of its noise.

### Mechanistic implementation of the *adder*

In order to have a biological interpretation for the division propensity, we provide a mechanistic framework for implementing the *adder* size control. Consider a timekeeping regulator that accumulates from zero molecules at the time of cell birth. The molecules of this regulator are assumed to be stable with a long half-life and synthesized according to a Poisson process with a rate *kf* (*s, τ*) proportional to the growth law. A special case of this is the exponential growth law *f* (*s, τ*) ∝*s*, where the division regulator is produced at a rate proportional to cell size [12, 28]. The theoretical foundation of this mechanism has been investigated in several previous works [25, 43, 44]. The number of molecules of our regulator *m*(*t*) ∈{0, 1, …} is a random process with integer values, with transitions from *m* − 1 to *m* molecules occurring stochastically with propensity *kf* (*s, τ*) (Figure 2A). From a biological perspective, the rate of change in cell size *f* (*s, τ*) corresponds to the global protein production rate. Therefore, the synthesis rate *kf* (*s, τ*) can be naturally interpreted as indicating that there is a fraction of proteome synthesis allocated to the expression of cell cycle regulators [45]. A cell division event is triggered when the cell cycle regulator accumulates to a threshold of *M*≥ 1 molecules, and the regulator is subsequently degraded during cytokinesis so that newborns restart the cycle with zero molecules.

**Figure 2:**
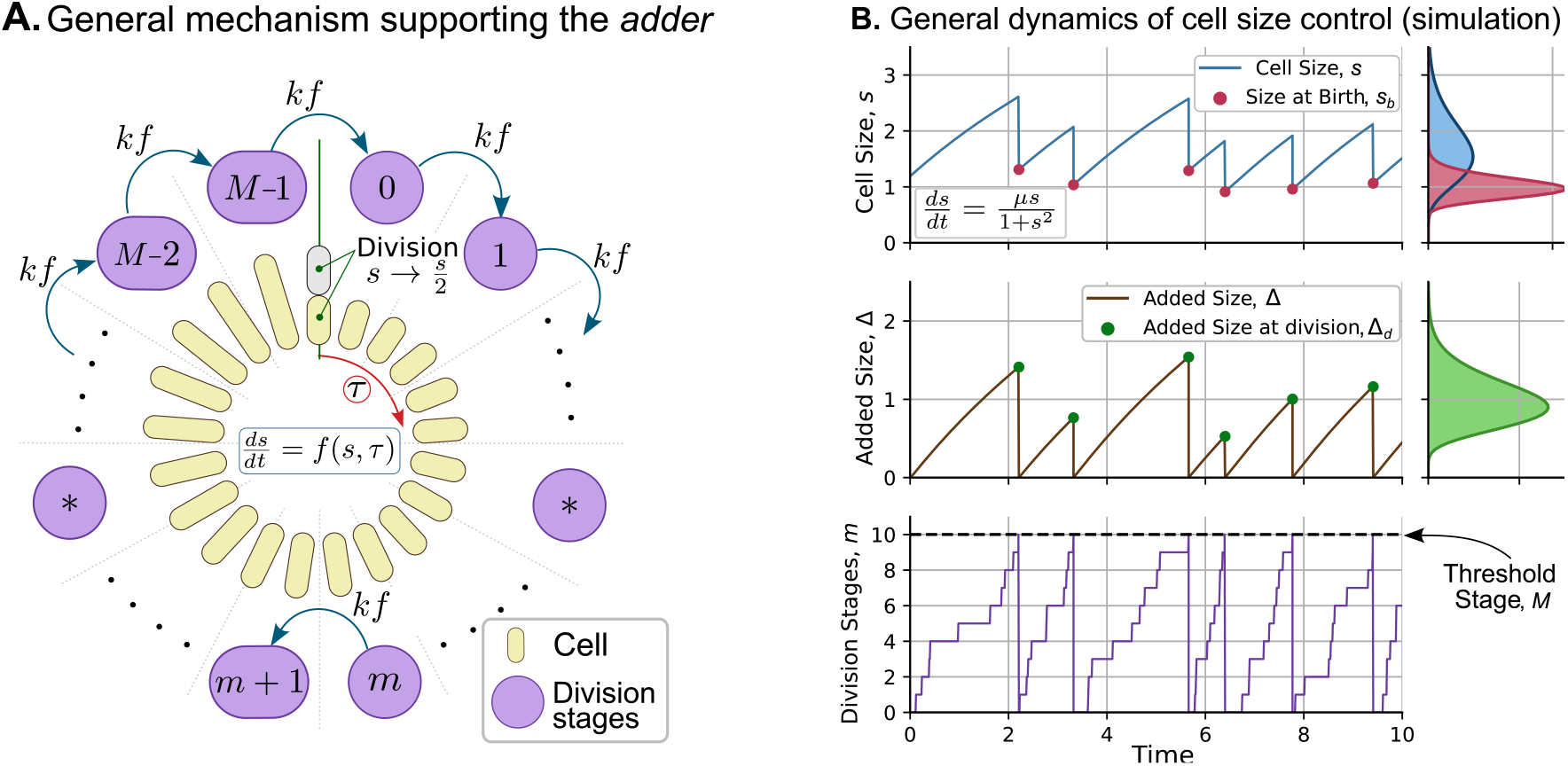
A general mechanism implementing *adder* based cell size homeostasis. **(A)** The cell size *s* grows according to (1) with an arbitrary growth law *f* (*s, τ*). During the cell cycle, cell performs transitions between different stages *m* ∈ {0, 1, …, *M*− 1} at a rate proportional to the growth law,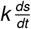. Upon reaching the *M*-th stage, the cell divides, and the newborns restart the cell cycle from stage *m* = 0. Alternatively, *m* can be interpreted as the number of molecules of a stable cell cycle regulator expressed at a rate 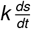, with cell division triggered upon reaching a critical threshold of *M* molecules. **(B)** Simulation of cell size over time for a general non-exponential growth law. *Top:* Cell size over time (solid line) showing different values of size at birth (red dots). The marginal plot shows their respective steady-state distribution. *Middle:* Added cell size as a function of time (solid line) showing different values of added size at division (green dots). The marginal plot shows the distribution of added size from cell birth to division. *Bottom:* Crossing of division stages *m*(*t*) over time. Once the cell reaches stage *M* = 10 (threshold stage), it undergoes cell division, and the stage is reset to *m* = 0. Simulation parameters: 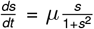; *µ* = 2.1, *k* = 10. The resets in cell size upon division were modeled as per (8) with *β* = 1*/*2 with probability one.

Alternatively, *M* can be interpreted as the number of stages a cell must pass through from birth until division. Newborns start with *m*(*t*_0_) = 0, and the transition between stages occurs stochastically with propensity *kf* (*s, τ*). In each transition from *m* − 1 to *m*, the added cell size is an exponentially distributed random variable with a mean of 1*/k*. Since there are *M >* 1 such transitions to reach the division threshold, the total added size during the cell cycle follows an Erlang distribution with a mean of ⟨Δ_*d*_⟩ = *M/k* and a squared coefficient of variation (Figure 2B):

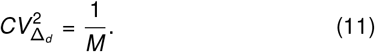

To connect this interpretation with our general theory, we observe that it is possible to obtain an associated propensity by estimating *h*(*y*) using the Erlang distribution as 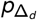 in (6). This mechanistic interpretation of the generalized adder has recently been applied in the analysis of cancer cells that were characterized by their higher 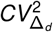 [36] compared to healthy ones.

This multi-step framework can be extended to different scenarios leading to an *adder* but with higher stochasticity in the added size. These scenarios include:

- The rates of transition from *m* to *m* + 1 can be generalized to be *k*_*m*_*f* (*s, τ*), *m*∈ *{*0, 1, 2, …, *M*−1}. In that case, the added size is a sum of *M* independent exponential random variables, each with mean 1*/k*_*m*_ [41].
- As previously done for the case of exponential growth [28], the expression of the regulator can occur in bursts, where bursts arrive as a non-homogeneous Poisson process with rate *kf* (*s, τ*), and each burst creates several molecules that are itself drawn from a size-invariant distribution.
- Finally, there could be statistical fluctuations in parameters *k* and *M* across cell cycles that correspond to some form of extrinsic noise in combination with the intrinsic noise of the regulator copy numbers.

Our proposed generalized model, while ensuring an *adder* regardless of the growth rate specifics, can be modified to capture other paradigms of cell size homeostasis, such as *timer* (cell cycle duration is independent of newborn size) or *sizer* (division size is independent of newborn size) [20, 46]. Such cases often result in an added size becoming correlated with the size at birth. For example, an adder-sizer combination would result in a negative correlation between Δ_*d*_ and *s*_*b*_ as in fission yeast [47] and slow-growing *E. Coli* cells [48]. This adder-sizer can be predicted by modifications to this mechanism, such as reversible steps in the stage-crossing process related to degradation of the division regulator [12, 48, 49]. Alternatively, this adder-sizer can also be modeled by accumlation of the regulator with a rate *ks*^*λ*^*kf* (*s, τ*) corresponding to size-dependent proteome allocation to regulator synthesis [46, 48, 50]. Here *λ >* 0 (*λ <* 0) captures the negative (positive) correlations between the added size and newborn size. Finally, the description of the division process using propensities has opened new perspectives in the characterization, not only of division strategies, but also of cell size dynamics across multiple cell species [46, 51, 52].

In summary, we have described both a phenomenological approach and a mechanistic approach implementing cell size homeostasis as per the *adder* principle irrespective of the growth law, and this approach can be easily generalized to other forms of cell size control.

### Clonal proliferation dynamics for different adders depends on the growth law

Next, we explore how cell size regulation at the single-cell level propagates across generations to affect the colony population. Can different types of growth laws yield qualitatively different fluctuations in clonal size, even though cell size control remains an *adder* ? To illustrate this, consider a cell colony that begins with a newborn cell (the progenitor) of size *s*_*b*_. Given that division events occur stochastically, the number of descendant cells *N*(*t*) from the single progenitor is a stochastic process. To characterize the random variability of the colony population size, we use its statistical moments: the mean number of cells in the colony ⟨*N*⟩ and the population noise quantified by 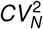, the squared coefficient of variation of *N*(*t*). We are specifically interested in connecting the transient dynamics of 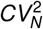 to the growth law *f* in for an *adder* with given statistical properties of Δ_*d*_.

### Modeling approach for computing 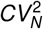

Statistical moments of *N*(*t*) can be approximated analytically [35] or numerically [53] in particular cases. To have a more accurate approach, in this article, we use individual-based simulation algorithms to simulate clonal expansion [34]. The stochastic timing of division is set by the requirement of the added size to division for each newborn cell to follow a prescribed distribution 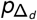, and each division event results in addition of two newborns to the population. To better illustrate the effect of the growth law on 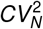, we simplify the approach by considering that all clonal replicas start with a single newborn with the same size, all cells follow the same growth law (i.e., no intercellular differences in growth rate). We further assume the added size to follow a gamma distribution with mean ⟨Δ_*d*_ ⟩ = 1 (arbitrary units) and noise in added size 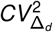. Note from (9a) that this choice of ⟨Δ_*d*_ ⟩ fixes the average size of the newborn cell to always be ⟨*s*_*d*_⟩ = 1. In this scenario, stochasticity in the added size is the only source of noise impacting clonal size dynamics and we modulate this noise level by considering two cases: a high noise 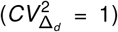 and a low .noise 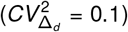 in Δ_*d*._

Referring the reader to supplementary section S2 for details on our simulation algorithm, we present sample trajectories of cell size, clonal population size, and total biomass (sum of cell sizes of all descendant cells at a given time point) in Figure 3 for exponential (top panel) and non-exponential growth in cell size (bottom panel).

**Figure 3:**
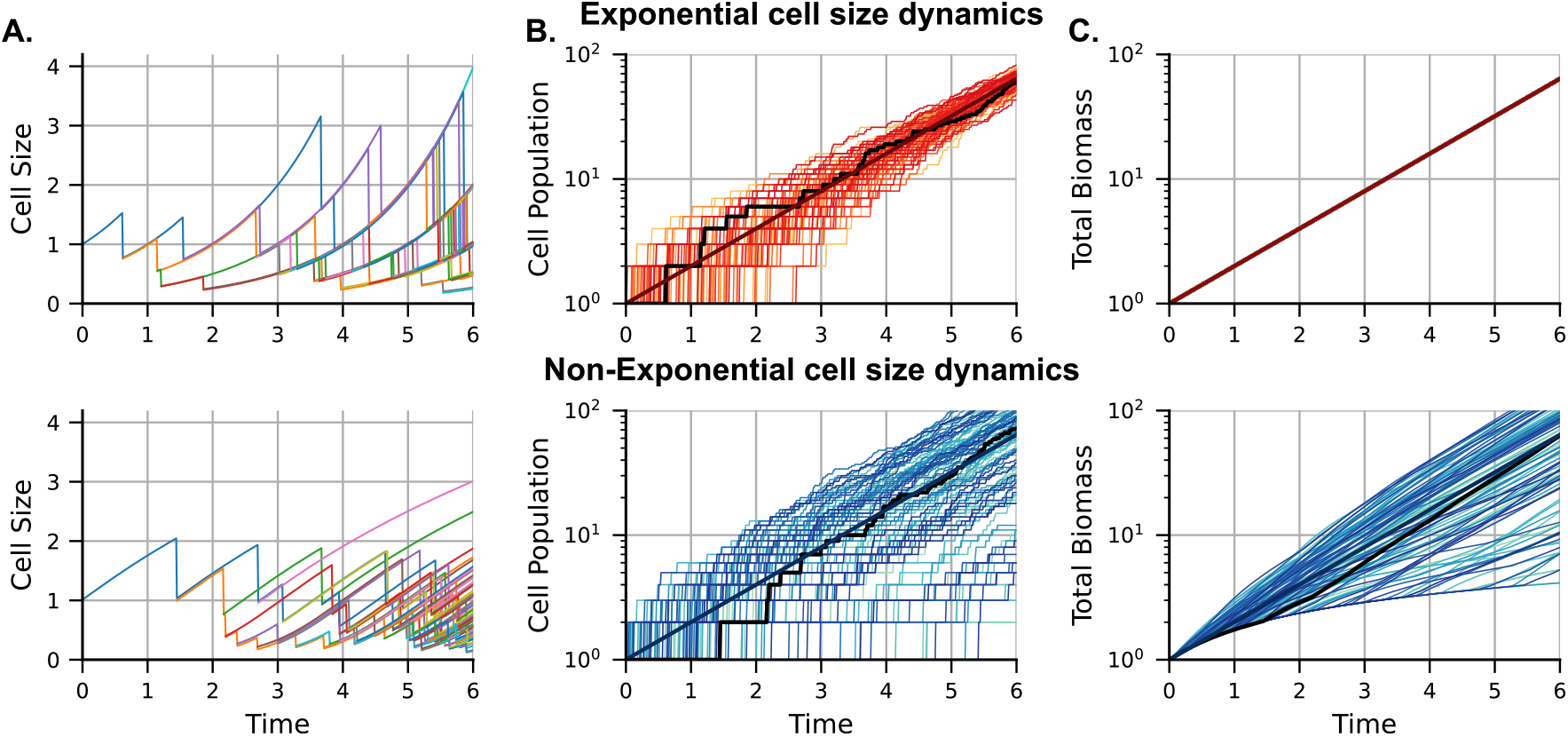
Illustration of sample trajectories for cell size, clonal population size, and total biomass for exponential and non-exponential cell size dynamics, with cell division timing set according to the *adder*. **(A)** (top panel) Cell size dynamics for exponentially 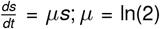) and (bottom panel) non-exponentially growing cells 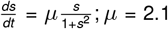 within a proliferating colony. Different colors represent different descendants of the colony progenitor. In both cases, size control follows the *adder* model, where the size added (Δ_*d*_) in each cell cycle is an independent and identically distributed random variable following an exponential distribution with ⟨Δ_*d*_ ⟩ = 1 and 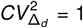. **(B)** Examples of clonal population size over time for simulated colonies. Each colony starts with a newborn progenitor cell with size *s*_*b*_ = 1 *a*.*u*. The black line represents the colony in **(A)**, and the dark red line is the mean population number. The y-axis is on a logarithmic scale. **(C)** Total colony biomass, defined as the sum of the cell sizes of all descendant cells at a given time. When all colonies start with the same-sized progenitor, the biomass is given by (13). For the exponential case (top panel), the total biomass does not vary across colony replicas, while for the nonexponential case (bottom panel), biomass shows considerable inter-colony differences.

As an example of non-exponential growth law, we use

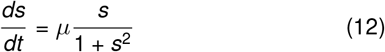

that was previously reported based on experiments measuring the proliferation potential of cell size-sorted human and *Drosophila* cells [54, 55].

### Exponential cell size dynamics

Our results show that for exponential cell size dynamics with *f*∝*s* in (1), 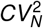 increases from zero reaching its maximum around the first division and then decreases monotonically to zero over time (Figure 4A). Thus, as the average clonal size expands exponentially (Figure 3B top), the interclonal population size differences asymptotically vanish. The difference between low-noise 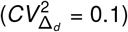 and high-noise 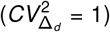 in added size is the transient behavior of 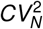, with 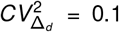 exhibiting a lower peak value of 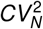 and oscillatory dynamics (Figure 4A).

**Figure 4:**
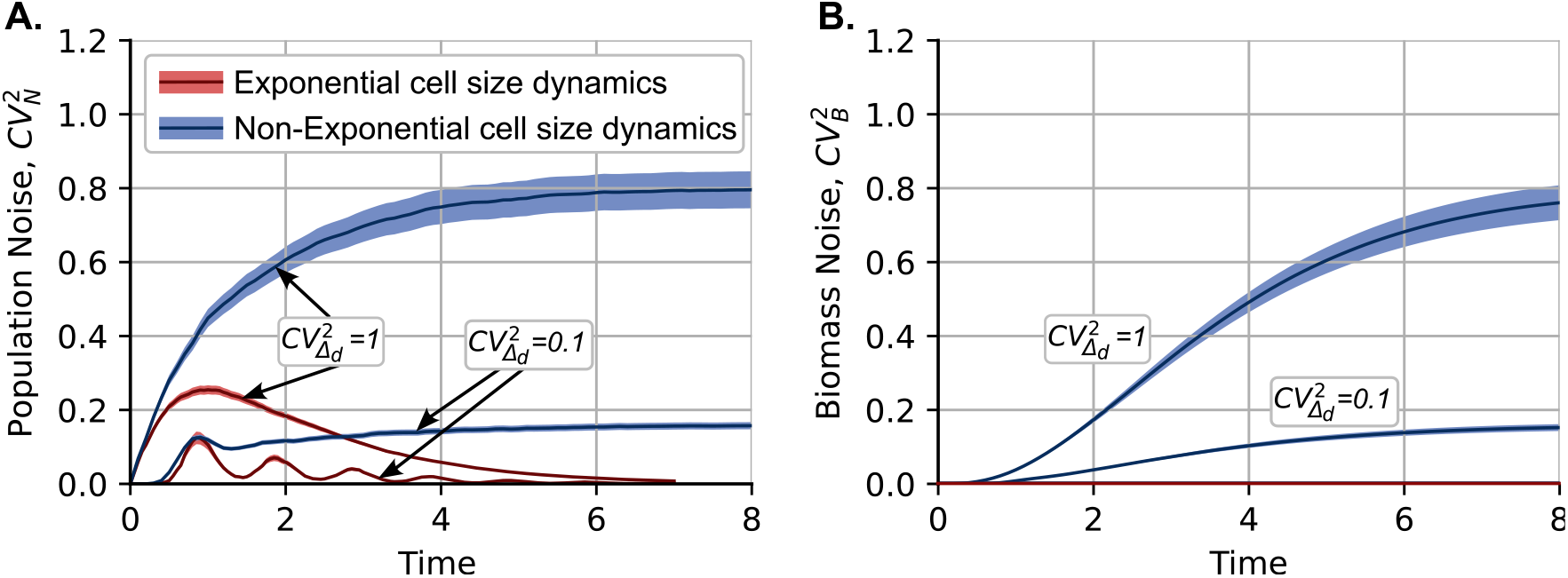
Cell size dynamics qualitatively impact the stochastic dynamics of clonal population size. **(A)** Plot of the squared coefficient of variation of clonal size 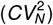 over time for cell size dynamics following exponential and non-exponential growth laws as described in Figure 3. In all cases, cell size homeostasis occurs according to the *adder* model, with Δ_*d*_ following a gamma distribution with ⟨Δ_*d*_ ⟩ = 1 and two values of 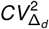 corresponding to high 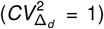 and low noise 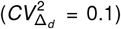. For exponential cell size dynamics, *CV*_*N*_ exhibits a transient peak and then approaches zero as *t* → ∞. In stark contrast, 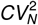 increases over time to asymptotically approach a non-zero value for non-exponential cell size dynamics. Lowering the noise in the added size 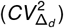 results in lower values of *CV*_*N*_. **(B)** Plot of the squared coefficient of variation 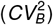 of total biomass, i.e., the aggregated size of all cells in a colony. For the exponential case, *CV*_*B*_ = 0 due to (13), while *CV*_*B*_ monotonically increases over time in the non-exponential case, exhibiting similar asymptotic values to *CV*_*N*_. All coefficients of variation are computed based on 5000 colony replicas, with colored regions representing the 95% confidence interval obtained using bootstrapping.

To intuitively understand why 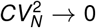 as *t* → ∞ it might be useful to consider the net biomass of the colony *B*(*t*) which is the aggregate size of all cells in a colony at a given time. For exponential cell size dynamics, the total biomass follows the simple expression

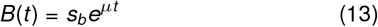

where *s*_*b*_ is the cell size of the progenitor cell that is assumed to be the same across colony replicas. Thus, despite the fact that the number of cells is variable from colony to colony, the total biomass is fixed between colonies at any time *t* with the coefficient of variation 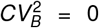. When the number of cells in the colony is very large (each having random differences in cell size), by the central limit theorem one could approximate 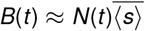, where 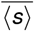 is the average steady-state cell size. From this approximation, it can be seen that

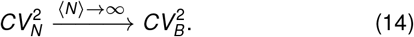

Note that for (14) to hold, the noise in cell size has to be bounded 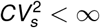. This property is ensured by cell size homeostasis mechanisms. A particular case where (14) does not hold is the timer-based cell division. In this mechanism, cell cycle duration is a size-independent random variable. Such timer-based are known to be non-homeostatic with variance in cell size increasing unboundedly over time [56].

### Non-exponential cell size dynamics

Interestingly, a qualitatively different result emerges when cell size dynamics follow non-exponential dynamics. This stark contrast is illustrated in Figure 3C. Unlike the exponential case, where biomass is given by (13) with no interclonal variability 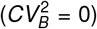, the non-exponential case shows biomass differences between clones due to the history of cell division events in the colony 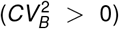. The variability in biomass across colony replicas is quantified in Figure 4B, where the biomass coefficient of variation (*CV*_*B*_) increases monotonically over time, eventually reaching a positive value. Tighter cell size regulation results in lower *CV*_*B*_ values (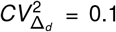compared to 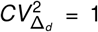). Consequently, as indicated by (14), since 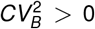, the population noise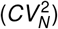) does not decay to zero but instead reaches a positive value after several generations (Figure 4A). Although 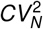 and 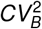 are both zero at the start of clonal expansion, simulations confirm that lim 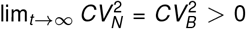. However, as shown in Figure 4, the transient profiles of these noises 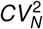 and 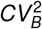 differ. Additional examples of the effect of non-linearity in the growth law on clonal size noise are provided in the supplementary section S3.

## Discussion

An important contribution of this article is to formulate a systems-level stochastic model implementing *adder* -based cell size homeostasis for any arbitrary cell size dynamics. This is achieved by appropriately defining a stochastic propensity of cell division events that ensures that the added size from cell birth to division is a random variable independent of newborn size and follows a given distribution 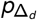. At any instant in the cell cycle, this propensity is given as the product of two functions: the growth law *f* in (1); and a function *h* defined in (6) of the added size since birth to that cell cycle time instant. We also proposed a mechanistic implementation of this scheme, which consists of a cell division regulator synthesized at a rate proportional to the cell size growth law, and division is triggered upon reaching a prescribed threshold level (Figure 2). In the context of bacteria, a potential regulator could be the FtsZ protein that polymerizes to form a ring (Z-ring) defining the site of division and recruiting other division proteins to the site [12]. The generalization of the *adder* presented here is particularly relevant given recent findings of an *adder* in several human cell lines [26, 57] despite exhibiting non-exponential cell size dynamics (see Figure 4 in [26]).

We connect *adder* -based cell size control at the single-cell level to variations in clonal size at the population level, as quantified by the coefficient of variation *CV*_*N*_ of cell number across colony replicates. The main finding is that despite having exactly similar *adder* mechanisms, the growth law *f* qualitatively changes the stochastic dynamics of clonal population expansion. For exponential cell size dynamics, our results show a transient peak in *CV*_*N*_ that decays to zero over time (Figure 4A), while for the non-exponential case, *CV*_*N*_ increases over time to reach a positive steady-state limit (Figure 4B). These qualitative differences can be explained by looking at the dynamics of the total colony biomass. Exponentially growing cells show smooth exponential growth of biomass that increases as in (13) and is invariant of the timing specifics of individual division events. In contrast, cells with non-exponential growth yield a colony biomass that depends on the history of division events (Figure 3C). The latter non-exponential scenario results in colony-to-colony biomass fluctuations that increase over time (Figure 4B), resulting in higher levels of *CV*_*N*_ approaching a non-zero steady state value as per (14).

An important limitation of the results presented in Figures 3-4 is that we considered the timing of cell division events as the only source of noise in the colony expansion process. Several biologically realistic factors, such as variability in the size of the initial progenitor cell and growth rate fluctuations, were ignored. In supplementary section S4, we repeated simulations of clonal expansion with these additional noise sources and, as expected, we observe that they amplify both *CV*_*N*_ and *CV*_*B*_ (Figure S2).

These findings have important implications for understanding the clonal expansion of cancer cells, for example, measured through barcoding technologies. More specifically, each cell receives a unique barcode at an initial time, and after several generations of clonal expansion, the number of cells in each barcode (or clone) is determined. Taking data from a recent publication on a melanoma cell line [58], we find a large variability in clonal population size at the endpoint (*CV*_*N*_≈1.3). Given that many human cell lines exhibit non-exponential cell size dynamics [26], this *CV*_*N*_ value is consistent with our results in Figure 4, where *CV*_*N*_ approaches a positive steady state. However, given the large variability seen in the data, it may be necessary to include other noise mechanisms (such as differences in the growth rate of single cancer cells within the same isogenic population) to capture the experimentally observed 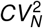. In contrast, cells with tighter cell size regulation and exponential growth, as seen with some bacterial species, yield lesser inter-clonal population size fluctuations [34]. Our study motivates time-resolved measurements of *CV*_*N*_ to systematically discern the noise sources driving clonal expansion differences, and such differences have important medical consequences driving single-cell heterogeneity in response to targeted cancer drug therapy [58–62].

## Author Contributions

- **Conceptualization:** C.N., C.A.V.G., A.S.
- **Investigation:** C.N.
- **Data Curation:** C.N.
- **Formal Analysis:** C.N., C.A.V.G., A.S.
- **Funding Acquisition:** A.S.
- **Methodology:** C.A.V.G., A.S.
- **Resources:** A.S.
- **Software:** C.N.
- **Supervision:** C.A.V.G., A.S.
- **Visualization:** C.N., C.A.V.G., A.S.
- **Writing:** C.N., A.S.
- **Writing – Review Editing:** C.N., C.A.V.G., A.S.

## Data accessibility statement

The simulation code used for these plots is publicly available at https://doi.org/10.5281/zenodo.13251255 [63].

## Acknowledgments

AS acknowledges the support of NIH-NIGMS via grant R35GM148351.

## Artificial intelligence

The authors declare that they did not use generative AI and AI-assisted technologies in the writing process.

