## supplementary Information for "A Generalized *Adder* mechanism for Cell Size Homeostasis: Implications for Stochastic Dynamics of Clonal Proliferation"

### S1 Proof for the Generalized Adder

Consider a cell born at time $t = t_0$ with initial size $s(t_0) = s_b$. Throughout the cell cycle, the monotonically increasing cell size $s(t)$ is given as the solution to (1). The cell division event occurs, at any point in the cell cycle, stochastically with a propensity given by $f(s, t-t_0)h(\Delta(t-t_0))$. Consequently, the cell-cycle duration $\tau_d$ is a random variable with the cumulative density function (CDF) [1] satisfying:

$$\mathbb{P}(\tau_d < x|s_b) = 1 - e^{-\int_{t_0}^{t_0+x} f(s,t-t_0)h(\Delta(t-t_0))dt}. \tag{S1}$$

Using the transformation $z = \Delta(t - t_0) = s(t) - s(t_0)$, it follows that

$$\frac{dz}{dt} = \frac{ds}{dt} = f(s, t - t_0). \tag{S2}$$

Using $\Delta(0) = 0$, (S1) can be rewritten as

$$\mathbb{P}(\tau_d < x|s_b) = 1 - e^{-\int_0^{\Delta(x)} h(z)dz}. \tag{S3}$$

Since $\Delta(\tau)$ is a strictly monotonically increasing function of $\tau$,

$$\mathbb{P}(\Delta_d < \Delta(x)|s_b) = \mathbb{P}(\Delta(\tau_d) < \Delta(x)|s_b) = \mathbb{P}(\tau_d < x|s_b), \tag{S4}$$

which, from (S3), implies

$$\mathbb{P}(\Delta_d < \Delta(x)|s_b) = 1 - e^{-\int_0^{\Delta(x)} h(z)dz}. \tag{S5}$$

This results in the CDF

$$P_{\Delta_d}(y) = \mathbb{P}(\Delta_d < y|s_b) = 1 - e^{-\int_0^{y} h(z)dz}, \tag{S6}$$

with the probability density function (PDF)

$$p_{\Delta_d}(y) = \frac{dP_{\Delta_d}(y)}{d\Delta_d} = h(y)e^{-\int_0^{y} h(z)dz}. \tag{S7}$$

**Data:** $CV^2_{\Delta_d}$, $\langle\Delta_d\rangle$, $t_{max} > 0$, $s_b$, $\mu$
**Result:** $\{times = [\tau_d^1, \tau_d^2, \cdots, \tau_d^N]$, $sizes = [s^1, s^2, \cdots, s^N]\}$
$\Delta_d^1 \sim p_{\Delta_d}$;
$s^1 = s_b$;
$\tau_d^1 = \{\tau|\Delta(\tau) = \Delta_d^1\}$;
$times = [\tau_d^1]$;
$sizes = [s^1]$;
$\tau_{min} = \tau_d^1$;
$\tau = \tau_{min}$;
**if** $\tau < t_{max}$ **then**
  **while** $\tau < t_{max}$ **do**
    $i^* = argmin(times)$;
    $\tau_{min} = min(times)$;
    **for** $i = 0;\ i < length(times);\ i = i + 1$ **do**
      $times[i] = times[i] - \tau_{min}$;
      $sizes[i] = sizes[i] + \int_0^{\tau_{min}} \frac{ds^i}{dt}dt$;
    **end**
    $sizes[i^*] = sizes[i^*]/2$;
    $\Delta_d^{i^*} \sim p_{\Delta_d}$;
    $times[i^*] = \{\tau|\Delta(\tau) = \Delta_d^{i^*}\}$;
    $N = length(times)$;
    $\Delta_d^{N+1} \sim p_{\Delta_d}$;
    $sizes[N + 1] = sizes[i^*]$;
    $times[N + 1] = \{\tau|\Delta(\tau) = \Delta_d^{N+1}\}$;
    $i^* = argmin(times)$;
    $\tau_{min} = min(times)$;
    $\tau = \tau + \tau_{min}$;
  **end**
**end**
**for** $i = 0;\ i < length(times);\ i = i + 1$ **do**
  $times[i] = times[i] - (t_{max} - (\tau - \tau_{min}))$;
  $sizes[i] = sizes[i] + \int_0^{t_{max}-(\tau-\tau_{min})} \frac{ds^i}{dt}dt$;
**end**

**Algorithm 1:** The algorithm simulates the proliferation of a single colony using the *adder* division strategy. It takes into account the mean added size, $\langle\Delta_d\rangle$, the variability of this size, represented by $CV^2_{\Delta_d}$, and the maximum simulation time, $t_{max}$. The algorithm calculates the array *times*, which indicates the time remaining until the next division for all cells in the colony. Simultaneously, it generates the array *sizes*, which contains the sizes at the time of division. The population is estimated by counting the number of elements in the *times* array. $\Delta_d$ is an independent identically gamma-distributed with its parameters detailed in equation (S9).

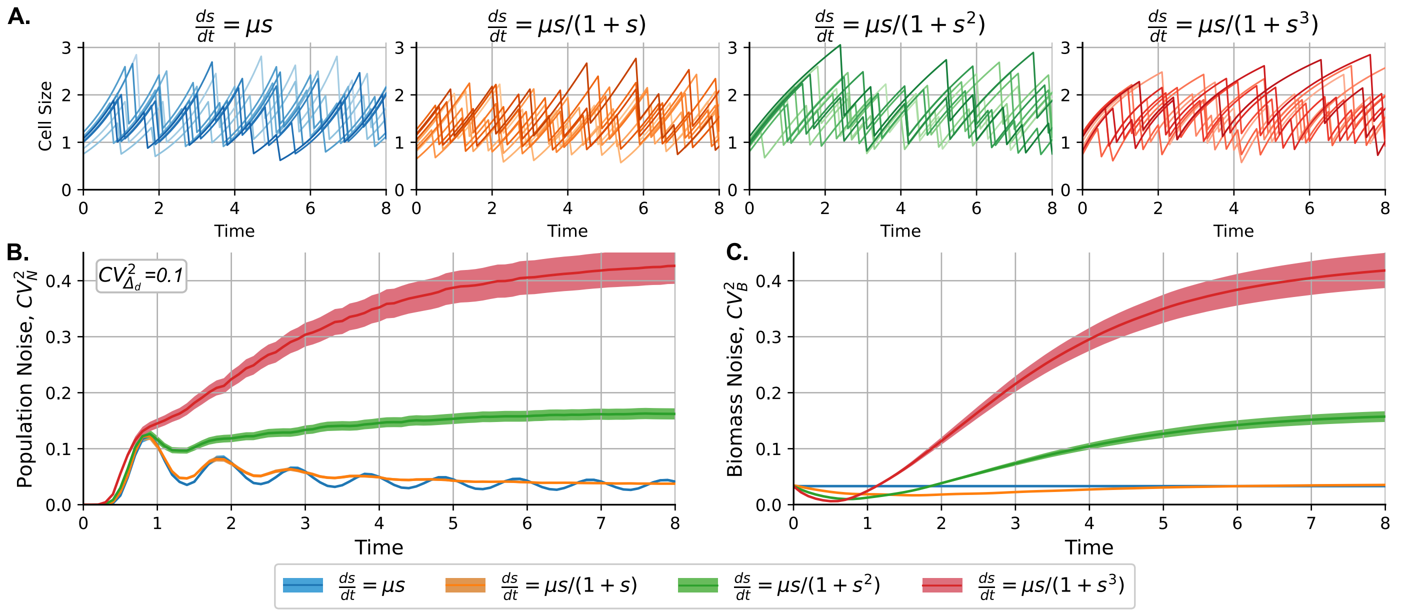


Figure S1: **Dynamics of population noise for different nonlinearities in the cell size growth law. (A)** Examples of cell size trajectories for different growth laws. We consider four cases of $\frac{ds}{dt} = \mu \frac{s}{1+s^{\alpha}}$, with $\alpha \in \{0, 1, 2, 3\}$. **(B)** Noise in cell population ($CV^2_N$) vs. time ($t$) for $CV^2_{\Delta_d} = 0.1$ and different nonlinear growth laws explained in **(A)**. **(C)** Noise in cell biomass ($CV^2_B$) vs. time ($t$) for $CV^2_{\Delta_d} = 0.1$ and different nonlinear growth laws. Parameters: $\langle s_b \rangle = \langle \Delta_d \rangle = 1$, $CV^2_{s_b} = \frac{CV^2_{\Delta_d}}{3}$ according to (10) since $CV^2_\beta = 0$. $\Delta_d$ follows a gamma distribution. All cells have the same $\mu$, chosen depending on $\alpha$ such that $\langle N \rangle$ doubles every unit of time. Division is considered to be perfectly symmetric with probability one ($\langle \beta \rangle = 0.5$, $CV^2_\beta = 0$).

### S2 Simulation details for the cell proliferation using the *adder* division

In this section, we present the algorithm used to simulate cell proliferation according to the *adder* division strategy. The pseudocode for this simulation is provided in Algorithm 1. The key parameters involved include the growth rate, denoted by $\mu$ (which represents the growth rate for exponential increases in cell size), the progenitor cell size $s_b$, the maximum simulation time $t_{max}$, the mean added size $\langle \Delta_d \rangle$, and its associated noise $CV^2_{\Delta_d}$.

An important input for the simulation is the progenitor cell size $s_b$, which follows similar statistics to any other newborn cell and is characterized by a size variability $CV^2_{s_b}$ and a mean size $\langle s_b \rangle$. We set $\langle s_b \rangle$ and $\langle \Delta_d \rangle$ to a value of 1.

To focus our simulation on the implications of the adder division strategy without delving into the specifics of the underlying mechanism, we parameterize the simulation using the added size statistics: the mean $\langle \Delta_d \rangle$ and the noise $CV^2_{\Delta_d}$. We assume that the added size, $\Delta_d$, follows a gamma distribution:

$$p_{\Delta_d}(y) = \frac{1}{\Gamma(\gamma)\theta} \left(\frac{y}{\theta}\right)^{\gamma-1} e^{-\frac{y}{\theta}}, \qquad \text{(S8)}$$

where $\Gamma(\gamma)$ is the gamma function, and the shape $\gamma$ and scale $\theta$ parameters are estimated from $\langle \Delta_d \rangle$ and $CV^2_{\Delta_d}$ using the following equations:

$$\gamma = \frac{1}{CV^2_{\Delta_d}}, \quad \theta = \langle \Delta_d \rangle \cdot CV^2_{\Delta_d}. \qquad \text{(S9)}$$

Given $\Delta_d$ for a particular cell cycle, we estimate the cycle duration $\tau_d$ based on the growth law. This is done from the size at birth, $s_b$, and the growth law. Once the added size evolves according to the dynamics described by equation (3) and $\frac{ds}{dt} > 0$, the cycle duration can be uniquely determined as the solution to $\tau_d := \{\tau | \Delta(\tau) = \Delta_d\}$, where $\Delta(\tau)$ is defined in (2).

For example, if we assume that cells grow at an exponential rate $\mu$, the division size $s_d$ follows the relationship $s_d = s_b e^{\mu \tau_d}$. In terms of the added size $\Delta_d$, this relationship can also be expressed as $s_d = s_b + \Delta_d$. Consequently, cell cycle duration $\tau_d$ can be determined in terms of $\Delta_d$ and $s_b$ as:

$$\tau_d := \{\tau | \Delta(\tau) = \Delta_d\} = \frac{1}{\mu} \ln\left(\frac{s_b + \Delta_d}{s_b}\right). \qquad \text{(S10)}$$

For more complex growth laws, this equation would need to be solved numerically.

The durations of all $N$ cell cycles, denoted as $\tau^i_d$ for each cell $i \in \{1, \ldots, N\}$, are stored in an array called *times*. Similarly, all cell sizes at division, $s^i_d$, are stored in an array called *sizes*. During each iteration, the minimum cycle duration, $\tau_{min}$, corresponding to a specific cell $i^*$, is identified. This $\tau_{min}$ is then subtracted from all $\tau^i_d$, and a new cell is generated with a birth size of $s^{i^*}_d/2$. The new cell is assigned a label $N+1$.

After generating the corresponding added size $\Delta^{N+1}_d$, the cell cycle duration $\tau^{N+1}_d$ is estimated and stored in the *times* array. A similar process of cell division and cycle duration estimation is performed for the original cell $i^*$ that just divided. The simulation time is then incremented by $\tau_{min}$, and the process repeats until the time to the next division exceeds $t_{max}$. At that point, the remaining time to $t_{max}$ is subtracted from all times to division. Finally, the number of cells is determined by the length of the *times* array.

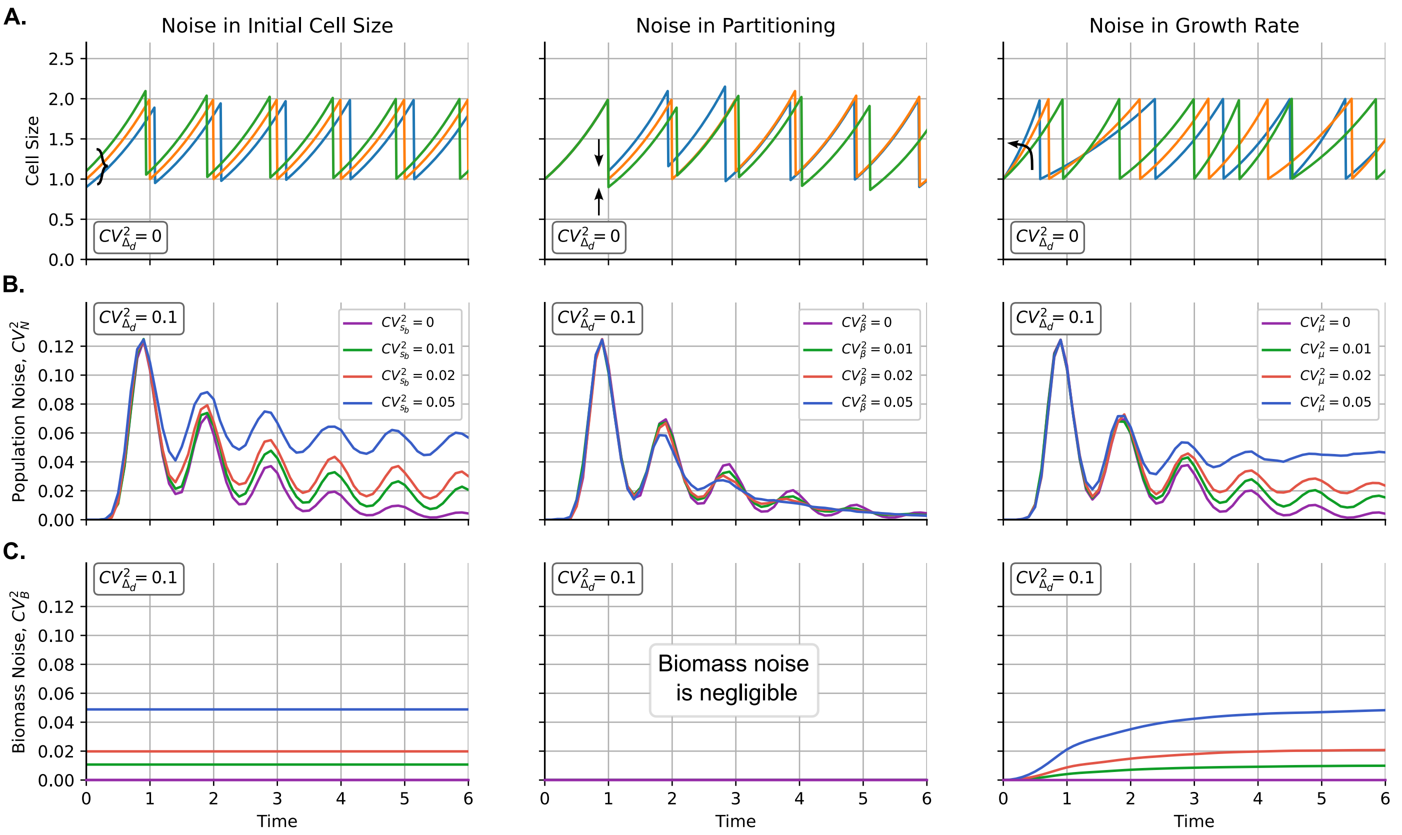


Figure S2: **Effect of different noise sources and cell size control on the dynamics of clonal size variability**. **(A)** Examples of cell size dynamics considering different sources of noise. For better illustration, the added size is deterministic $\langle \Delta_d \rangle = 1$, $CV^2_{\Delta_d} = 0$. Different colors represent different cells neglegting their descendants. **(B)** Dynamics of the variability of the population number $CV^2_N$ vs. time ($t$) for different sources of noise and $CV^2_{\Delta_d} = 0.1$. **(C)** Variability in biomass $CV^2_B$ vs. time ($t$) for different sources of noise and $CV^2_{\Delta_d} = 0.1$. (left) Dynamics considering different noise in progenitor cell size $CV^2_{s_b}$. (center) Dynamics for different noise in cell partioning $CV^2_{\beta}$. (right) Dynamics considering different noise in progenitor cell size $CV^2_{\mu}$. Parameters: $\langle \mu \rangle = \ln(2)$, $\langle s_b \rangle = \langle \Delta_d \rangle = 1$, $\langle \beta \rangle = 0.5$. The progenitor size ($s_b$) is gamma-distributed with specific noise ($CV^2_{s_b}$). $\beta$ is beta-distributed with $\langle \beta \rangle = 0.5$ and specific noise ($CV^2_{\beta}$). The growth rate ($\mu$) is gamma-distributed with $\langle \mu \rangle = \ln(2)$ and specific noise ($CV^2_{\mu}$).

### S3 Effects of a non linear growth law on population randomness

In the main article, we compare the dynamics of $CV^2_N$ for two examples of different growth laws: $\mu s$ and $\frac{\mu s}{1+s^2}$. Here, we extend our analysis by using our simulator to estimate colony dynamics under additional growth laws. To explore a range of representative cases, we consider the function $\frac{ds}{dt} = \frac{\mu s}{1+s^{\alpha}}$. This function allows us to model various growth behaviors:

- For $\alpha = 0$, we obtain exponential growth.
- For $\alpha = 1$, the model represents exponential growth that saturates to linear growth as cells become larger, a behavior recently observed in some bacteria [2].
- $\alpha = 2$ was previously reported in a heuristic model [3].
- For $\alpha = 3$, the growth rate decreases more rapidly with increasing cell size, which can be used to model organisms such as yeast, where the growth of the mother cell slows significantly after reaching maturation.

Figure S1A shows examples of cell size dynamics for each selected growth scenario. The initial cell size, $s_b$, is treated as a random variable with statistics $\langle s_b \rangle = \langle \Delta_d \rangle$ and noise $CV^2_{s_b}$, which is related to the noise in added size according to equation (9b), while ignoring partitioning effects:

$$CV^2_{s_b} = \frac{1}{3} CV^2_{\Delta_d}. \tag{S11}$$

The dynamics of $CV^2_N$ are illustrated in Figure S1B, where it is evident that the behavior of $CV^2_N$ is similar across different growth laws before the first division, but diverges significantly in subsequent generations. Specifically, exponential growth leads to robust oscillations, whereas other growth scenarios exhibit damped dynamics. A comparison between these dynamics and the dynamics of biomass noise, $CV^2_B$, is shown in Figure S1C. As described in the main text, the noise in biomass exhibits an asymptotic behavior similar to the noise in population number, though with distinct transient dynamics. Interestingly, under non-linear growth conditions, $CV^2_B$ initially decreases during the first generation but then increases rapidly in subsequent generations.

### S4 Effects of different noise sources on the dynamics of $CV^2_{\Delta_d}$

In the main article, we examine two primary sources of variability in cell colonies that grow and divide according to the *adder* model. The first source of variability is the noise in cell cycle duration, $\tau_d$. This noise can be adjusted by considering multiple division steps or by modeling the division propensity. The second source of variability is the non-linearity of the growth law, $ds/dt$.

In this section, we explore how additional sources of noise impact overall variability in the colony population, quantified as $CV_N^2$. Specifically, we consider the noise in progenitor cell size, partitioning noise, and growth rate noise. For simplicity, we present examples where the growth law is proportional to cell size, indicating exponential growth. The results can be qualitatively extended to non-exponential growth.

**Noise in the initial progenitor cell size**

Let $s_b$ be the size of the colony's ancestor cell at $t = 0$. For simplicity, Figure 3 assumes that this progenitor cell begins its dynamics with a size $s_b = \langle \Delta_d \rangle = 1$ with probability one. However, experimentally, $s_b$ exhibits variability that approximately follows the expression (S11) [4,5].

We can extend our analysis to the more general case where $s_b$ has any degree of noise. To understand how population growth dynamics are influenced by variability in progenitor size, consider that the progenitor cell size $s_b$ at $t = 0$ is now treated as a random variable. The progenitor will grow exponentially at a rate $\mu$. We define the sequence of division event times as $t_1, t_2, \ldots, t_J$, where $0 < t_1 < t_2 < \cdots < t_J < t$, measured from the beginning of the experiment, with $J \in \mathbb{N}$ representing the total number of division events before time $t$. From this sequence, we can show that the size of a cell at time $t$ is given by [6]:

$$s(t; J, s_b) = s_b \frac{\exp(\mu t)}{2^J}. \qquad \text{(S12)}$$

Figure S2A (left) illustrates several cell size trajectories with different initial sizes. To simplify the concept, the trajectories shown in Figure S2A (left) assume a deterministic added size with $CV_{\Delta_d}^2 = 0$. However, this explanation can be extended to the general case where $CV_{\Delta_d}^2 > 0$. Notice that the primary effect of a noisy progenitor cell size is a variation in the phase of the cells. This means that while the cells exhibit similar periodic dynamics, the time of division varies, maintaining approximately the same interval between divisions for cells with ancestors of different sizes. Specifically, smaller progenitors typically experience a longer delay before their first division compared to larger progenitors. After this initial division, the cell population grows asymptotically at an exponential rate. As a result, colonies derived from smaller cells generally have a smaller population than those derived from larger cells.

We consider the size of the progenitor cell to be drawn from a gamma distribution with a mean $\langle s_b \rangle$ and random variability quantified by the squared coefficient of variation, $CV_{s_b}^2$. A key consequence of increasing $CV_{s_b}^2$ is an amplification of noise in population fluctuations. Specifically, $CV_N^2$ does not converge to zero, but instead oscillates and stabilizes at a finite value close to $CV_{s_b}^2$ as $t$ approaches infinity (Figure S2B, left). In contrast, biomass variability does not change over time, as biomass in all colonies grows deterministically according to equation (13). The corresponding biomass noise is illustrated in Figure S2C, left.

**Noise in cell size partitioning**

Another source of noise that can influence the dynamics of $CV_N^2$ is the variability in size partitioning, where cells are not divided exactly in half. As illustrated in Figure 1A (center), we can simulate this by multiplying the cell size by a random variable $\beta$ after each division, where $\langle \beta \rangle = 0.5$ and the variability is quantified by $CV_\beta^2 > 0$. This implies that the size of a particular cell after $J$ divisions follows:

$$s(t; J, s_b) = s_b \exp(\mu t) \prod_{j=1}^{J} \beta_j, \qquad \text{(S13)}$$

where $\beta_j$ is the division parameter for the $j$-th division. In our simulation, we assume that $\beta$ follows a beta distribution with $\langle \beta \rangle = 0.5$ and the desired $CV_\beta^2$.

Although this partitioning noise affects cell size statistics, it does not alter the total biomass, and therefore the asymptotic value of $CV_N^2$ still approaches zero as $t \to \infty$ (Figure 1B, center). As $CV_\beta^2$ increases, the oscillations in $CV_N^2$ become more damped, but $CV_N^2$ continues to trend towards zero over time. The example trajectories in Figure 1A (center) show how cells lose their cell cycle syncronization over time after a division with partitioning noise. However, while one of the descendant cells may be smaller than expected due to noise, other cells tend to be larger. In general, the effects on each descendant cell seem to balance each other out, leaving the noise in the biomass negligible (Figure 1C, center).

**Noise in growth rate**

In general, cells in a population grow at different rates. Therefore, the observed population growth rate is an average. In the main text, for illustration, we assume that cells grow at a constant deterministic rate $\mu$ throughout all cell cycles. This deterministic growth assumption provides a baseline for our analysis. To incorporate stochasticity from various sources, we now consider the growth rate to be a random variable. Specifically, after each division $j$, a cell grows during that cycle at a rate $\mu_j$, where $\mu_j$ is an independent and identically distributed (i.i.d.) variable drawn from a gamma distribution centered on $\mu$ with variability quantified by $CV_\mu^2$. We neglect any correlations in growth rate between successive cycles [4,7,8] or correlations with other variables such as size at birth [7]. In the scenario of zero partitioning noise, the size $s$ at time $t$ after $J$ divisions, each occurring at time $t_j$, is given by:

$$s(t, J, s_b) = \frac{s_b}{2^J} \left( \prod_{j=0}^{J-1} \exp\left(\mu_j (t_{j+1} - t_j)\right) \right) \exp\left(\mu_J (t - t_J)\right). \qquad \text{(S14)}$$

Figure S2A (right) shows examples of cell size trajectories over time under this type of stochastic growth. Notice that not only do cells lose synchronization in their division times, but cells that grow more slowly also proliferate at a slower rate. The effect of this noise in growth rate on $CV_N^2$ is depicted in Figure S2B (right), where $CV_\mu^2$ is increased while keeping the mean added size $\langle \Delta_d \rangle$ constant. The main observation is that population fluctuations, $CV_N^2$, exhibit damped oscillations, and $CV_\mu^2$ influences the asymptotic limit of $CV_N^2$ as $t \to \infty$.

[5] César Nieto, Sarah Täuber, Luisa Blöbaum, Zahra Vahdat, Alexander Grünberger, and Abhyudai Singh. Coupling cell size regulation and proliferation dynamics of *C. glutamicum* reveals cell division based on surface area. *bioRxiv*, 2023.

[6] Cesar Augusto Nieto-Acuna, Cesar Augusto Vargas-Garcia, Abhyudai Singh, and Juan Manuel Pedraza. Efficient computation of stochastic cell-size transient dynamics. *BMC Bioinformatics*, 20(23):1–6, 2019.

[7] Maryam Kohram, Harsh Vashistha, Stanislas Leibler, BingKan Xue, and Hanna Salman. Bacterial growth control mechanisms inferred from multivariate statistical analysis of single-cell measurements. *Current Biology*, 31(5):955–964, 2021.

[8] Ping Wang, Lydia Robert, James Pelletier, Wei Lien Dang, Francois Taddei, Andrew Wright, and Suckjoon Jun. Robust growth of escherichia coli. *Current Biology*, 20(12):1099–1103, 2010.
